# Effects of pen enrichment on leg health of fast and slower-growing broiler chickens

**DOI:** 10.1101/2021.07.23.453550

**Authors:** Bahadır Can Güz, Ingrid C. de Jong, Carol Souza Da Silva, Fleur Veldkamp, Bas Kemp, Roos Molenaar, Henry van den Brand

## Abstract

Pen enrichment for broiler (meat-type) chickens is one of the potential strategies to stimulate locomotion and consequently contribute to leg health and welfare. This study was designed to evaluate effects of using a plethora of pen enrichments (barrier perches, angular ramps, horizontal platforms, large distance between feed and water and providing live Black Soldier fly larvae in a dustbathing area) on tibia characteristics, locomotion, leg health and home pen behaviour of fast and slower-growing broiler chickens. The experiment was set up as a 2 x 2 factorial arrangement with a total of 840 male broiler chickens in a complete randomized design (7 replicates per treatment and 30 chickens per replicate) with the following treatments: 1) pen enrichment (enriched pen or non-enriched pen); 2) broiler strain (fast-growing Ross 308 or slower-growing Hubbard JA 757). Home pen behaviour and use of enrichment were observed. At approximately 1400 and 2200 gram body weight, two chickens per pen were randomly selected and slaughtered, to investigate tibia morphological, biophysical and mechanical characteristics and leg health. Pen enrichment positively affected tibia biophysical characteristics, e.g., osseous volume (Δ=1.8 cm^3^, *P*=0.003), total volume (Δ=1.4 cm^3^, *P*=0.03) and volume fraction (Δ=0.02 %, *P*=0.002), in both fast and slower-growing chickens, suggesting that pen enrichment particularly affects ossification and mineralization mechanisms. Accordingly, locomotion and active behaviours were positively influenced by pen enrichment. However, pen enrichment resulted in lower body weight gain in both strains, which might be due to higher activity or lower feed intake as a result of difficulties of crossing the barrier perches. Regarding the strain, slower-growing chickens showed consistently more advanced tibia characteristics and more active behaviour than fast-growing chickens. It can be concluded that pen enrichment may lead to more activity and better bone development in both fast and slower-growing chickens.

## Introduction

In the last decades, genetic selection on growth rate and feed efficiency in broiler (meat-type) chickens resulted in significant phenotypic and genotypic changes [1,2,3,4]. Despite the fact that this selection has provided numerous advantages e.g., high amount of meat production in a short rearing duration, less environmental pollution and considerable financial benefits for producers, it has also caused some downsides e.g., suboptimal leg health. Suboptimal leg health appears to be related to an imbalance between high growth rate and immature bones and joints [2,5,6,7,8], which can lead to impaired locomotion [2,6,8,9], pain [8, 10], poor welfare [8,11,12,13], higher mortality, lower slaughter revenues and significant financial losses [2,14,15,16,17].

A potential strategy to promote leg health and welfare of modern broiler chickens might be to stimulate activity and locomotion, e.g., by pen enrichment [18,19,20,21]. Chickens have been using natural perches, platforms, ramps and elevated resting areas as their natural behaviour throughout their history, from wild ancestors to their modern generations [21,22,23]. This suggests that these types of enrichments are important to fulfil natural behaviors, but current broiler houses mostly lack any form of enrichment. Several studies assessing behaviour showed that broiler chickens spend approximately 80% of their lifespan with passive behaviours (e.g., lying, sitting and resting) [4,18,24]. The lack of activity, together with a fast growth rate, may impair bone development, which can result in suboptimal leg health or even lameness [12,18,25,26,27]. It has been shown that a lower stocking density [21,28,29,30,31], placing platforms and/or ramps [21,23,32,33], perches [31, 32], large distance between feed and water [18,33,34], different dustbathing materials, such as moss-peat [35], and worms or insects in a dustbathing area [36, 37] resulted in lower prevalence of leg disorders and lower mortality rate, although we did not find this in an earlier comparable study [38]. Increasing physical activity and locomotion may thus result in lower incidence of leg problems by stimulating tibia morphological, biophysical and mechanical properties [18,33,39,40,41].

Another potential strategy to promote leg health and welfare is to reduce growth rate of broiler chickens. Fast-growing broiler chickens demonstrate more leg and locomotion problems than slower-growing broiler chickens [14,42,43]. The underlying reason is that fast-growing broiler chickens have more porous and less mineralised bones than slower-growing broiler chickens, which are less able to carry the rapidly increased body weight [44, 45]. It has been found that slower-growing broiler chickens spent more time on perches and platforms [46, 47], demonstrated better locomotion [24,26,46,48,49,50], had less hock and leg problems [46, 51] and lower mortality [52] than fast-growing broiler chickens.

It can be hypothesized that pen enrichment positively affects bone development and locomotion in both fast and slower-growing broiler chickens, but that effects might be larger in the fast-growing broiler chickens, because they generally show less locomotion. However, effects of pen enrichment on locomotion and leg problems in slower-growing broiler chickens are hardly investigated.

The aim of this study was to investigate effects of a combination of different forms of pen enrichment on tibia characteristics, locomotion, leg health and home pen behaviour of both fast and slower-growing broiler chickens.

## Materials and methods

### Experimental Design

The experiment was setup as a 2 x 2 factorial arrangement with two strains of broiler chickens (fast-growing or slower-growing) and two different levels of pen enrichment (enriched or non-enriched). A total of 28 pens (7 pens per treatment, each containing 30 male broiler chickens) within a complete randomized design was used. Fast-growing broiler chickens were reared till day 38 of age, whereas slower-growing broiler chickens were reared till day 49 of age. The experiment was conducted at the research accommodation of Wageningen Bioveterinary Research (Lelystad, The Netherlands). All procedures in this study were approved by the Central Commission on Animal Experiments (The Hague, The Netherlands; approval number: 2016.D-0138.006).

### Animals, Rearing and Housing Management

A total of 420 fast-growing (Ross 308, breeder age of 30 weeks) and 420 slower-growing (Hubbard JA 757; breeder age of 28 weeks) day-old male broiler chickens were obtained from a commercial hatchery (Probroed, Groenlo, The Netherlands) and randomly allocated to 28 pens in one broiler house. Half of the chickens per broiler strain were placed in enriched pens, while the other half was placed in non-enriched pens, resulting in the following treatments: enriched fast (**EF**), non-enriched fast (**NF**), enriched slower (**ES**) and non-enriched slower (**NS**). Pen size of both enriched and non-enriched pens was 3 x 1 m and floors in all pens were covered with wood shavings as bedding material. Enriched pens contained two wooden platforms (100 x 20 x 40 cm, one at each long side of the pen), two wooden ramps (200 x 20 cm, angle of 11.5°), a dust bathing area in the centre of the pen (100 x 100 cm) with peat moss (with a thickness of 2 cm in week 1, 4 cm in week 2, and 7.5 cm from week 3 onwards), two vertical wooden barrier perches (100 x 4 cm, adjustable in height from 4 to 16 cm with steps of 4 cm at days 7, 14 and 21), a maximum distance (3 m) between feeders and drinkers and provision of live Black Soldier fly larvae (**BSFL**) in the substrate of the dust bathing area (once daily between 11:00 and 11:15 h). The amount of BSFL was determined daily, based on 5% of the expected feed intake, except during the first 7 days, where chickens received a higher level of BSFL (10% on days 0 – 1, 15% on days 2 – 4 and 10% on days 5 – 7). The reason for using higher percentages in these 7 days is related to the number of larvae available for each chicken. With the low feed intake in this phase, only one or two larvae would have been available per chicken in case only 5% BSFL was provided. Non-enriched pens included feed and water (at 1 m distance) and one single long perch (300 x 4 cm, not adjustable in height). Illustrations of the enriched and non-enriched pens are provided in Fig 1.

**Fig 1.**
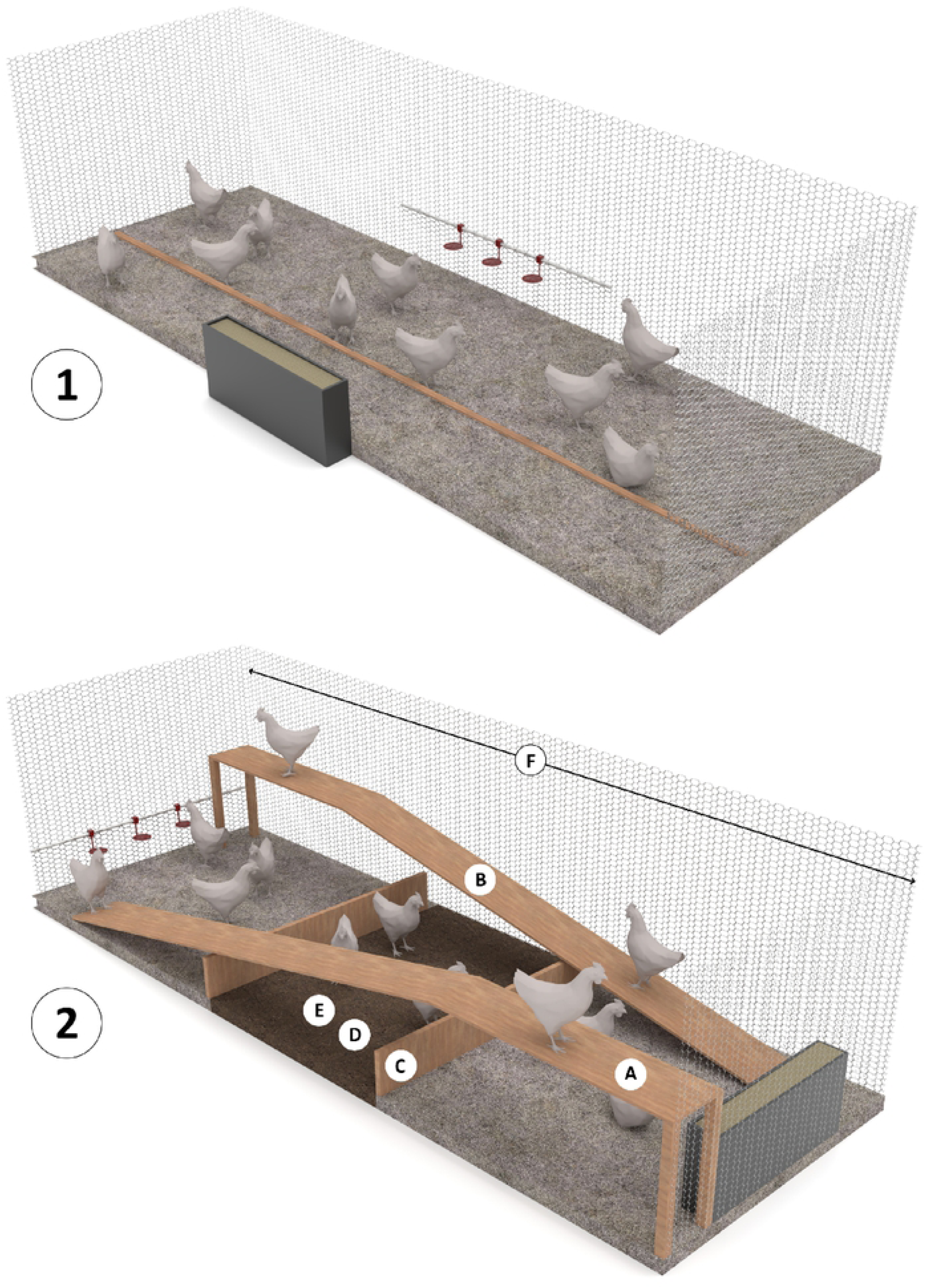
Illustrations of non-enriched (top) and enriched (bottom) pens. Non-enriched pens (3 x 1 m) contained a short distance (1 m) between feeders and drinkers placed on opposite long walls, had one non-adjustable perch and the pen was covered with wood shavings as a bedding material. Enriched pens (3 x 1 m) contained two wooden platforms (A; 100 x 20 x 40 cm, one at each long side of the pen), two wooden ramps (B; 200 x 20 cm, angle of 11.5°), two vertical wooden barrier perches (C; 100 x 4 cm, adjustable in height from 4-16 cm with steps of 4 cm at days 7, 14 and 21), dust bathing area (D; 100 x 100 cm) with peat moss (with depth of 2 cm in week 1, 4 cm in week 2, and cm from week 3 onward), provision of live Black Soldier fly larvae (BSFL) in the substrate of the dust bathing area (E; once daily between 11:00 and 11:15 h) and a large distance (3 m) between feeders and drinkers placed on opposite short walls (F). The floor outside the dust bathing area was covered with wood shavings as a bedding material.

At day 0 (placement), all broilers were provided with a neck tag for individual identification. House temperature was maintained at 34°C at day 0 and gradually decreased to a constant temperature of 18°C at 40 days of age. Relative humidity was kept between 60% and 80% from 1-7 days of age and between 40% and 60% thereafter. The lighting program used was 24L:0D (day 0), 20L:4D (day 1 to 6) and 18L:6D (from day 7 onward, with a continuous dark period during night). At day 0, chickens were vaccinated against infectious bronchitis (eye drop; MSD Animal Health, Boxmeer, The Netherlands) and at day 11, against Newcastle disease (Clone 30; eye drop, MSD Animal Health, Boxmeer, The Netherlands).

Feed and water were provided *ad libitum* for all treatments throughout the whole experiment. A 3-phase feeding program was applied; starter diets were provided from day 0 to 14 (ME=2925 kcal/kg, CP=203 g/kg, dLys=11.1 g/kg), grower diets from day 14 to 35 (ME=2975 kcal/kg, CP=171 g/kg, dLys=9.1 g/kg) and finisher diets from day 35 to 38 (for fast-growing chickens) or 35 to 49 (for slower-growing chickens) (ME=3025 kcal/kg, CP=165 g/kg, dLys=8.6 g/kg). Coccidiostats (70 g/kg salinomycin) were added to the grower diet. A protein-fat mixture, with a comparable composition as the BSFL, was added to the diet of the non-enriched pens once daily to achieve similar energy and nutrient intake as the broilers in the enriched pens (which received BSFL).

### Data Collection, Sampling and Measurements

All chickens were individually weighed on day 0, 7, 14, 21, 28, 35, 42, and 49 of age. Feed intake (**FI**) was determined per pen at the same days. Body weight (**BW**), FI and feed conversion ratio (**FCR**) were calculated for the three phases and over the whole growing period, taking mortality into account. FI was calculated without BSFL intake of chickens in enriched pens and by excluding the intake of the protein-fat mixture of chickens in non-enriched pens. Mortality was recorded per pen per day. Home pen behaviour (all chickens per pen) and use of enrichment (all chickens per pen in enriched pens) were scored by direct observation of one observer, using instantaneous scan sampling [53] at day 8, 22, 29 and 43. At these days, broilers were observed in their home pen at four moments (8:30, 10:30, 13:00 and 15:00 h). On day 43, only slower-growing chickens were present. Per scan per day per pen, behaviour of all chickens was scored during 3 to 4 min. The number of chickens performing the following behaviours was scored: eating, drinking, walking, standing, sitting, comfort behaviour, foraging, dustbathing, ground pecking, aggression and others. Others was defined as chickens demonstrating a behaviour other than all other behaviours described above. After observing the behaviour in a pen, the number of chickens performing the following activities was scored for use of enrichment: chickens on platforms and ramps, chickens under platforms and ramps, dustbathing chickens and chickens perching on barrier.

Gait score of 4 randomly selected chickens per pen was evaluated on day 27 (fast-growing chickens) and day 35 (slower-growing chickens), to eliminate BW difference. Gait was scored within a range of 0 (normal locomotion) to 5 (unable to walk) [54].

At day 29 and 38, two fast-growing chickens per pen were selected for slaughtering with an average body weight of 1400 and 2200 gram, respectively, whereas at day 38 and 49, two slower-growing chickens per pen were selected for slaughtering with the same body weights. Chickens were subjected to electrical stunning for euthanizing. Then, Varus-Valgus (**VV**) was scored, after fixating the legs at the hip joint to stretch the leg, by determining the angle between the tibia and the metatarsus for both the left (**VV^L^**) and right leg (**VV^R^**), using a goniometer. Thereafter, the left leg of each chicken was dissected and assessed by a veterinarian on tibia dyschondroplasia (**TD**), bacterial chondronecrosis with osteomyelitis (**BCO**), epiphyseal plate abnormalities (**EPA**), and epiphysiolysis (**EPI**). All these leg disorders were scored in the range of 0 (no abnormalities), 1 (minor abnormality) or 2 (severe abnormality).

Right legs were deboned and tibias were packed and frozen at -20°C. After thawing, tibia weight was determined. Tibia proximal length, lateral cortex thickness, femoral and metatarsal side proximal head thickness, osseous volume, pore volume, total volume (osseous volume + pore volume), volume fraction (osseous volume / total volume), mineral content and mineral density were analysed on individual tibia, using a GE Phoenix 3D X-ray microfocus CT scanner (General Electric Company^®^, Boston, Massachusetts, US); for details see [55, 56]. Illustrations of scanned bones are provided in Fig 2. Robusticity index was calculated using the following formula [57]:

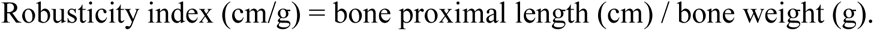

**Fig 2.**
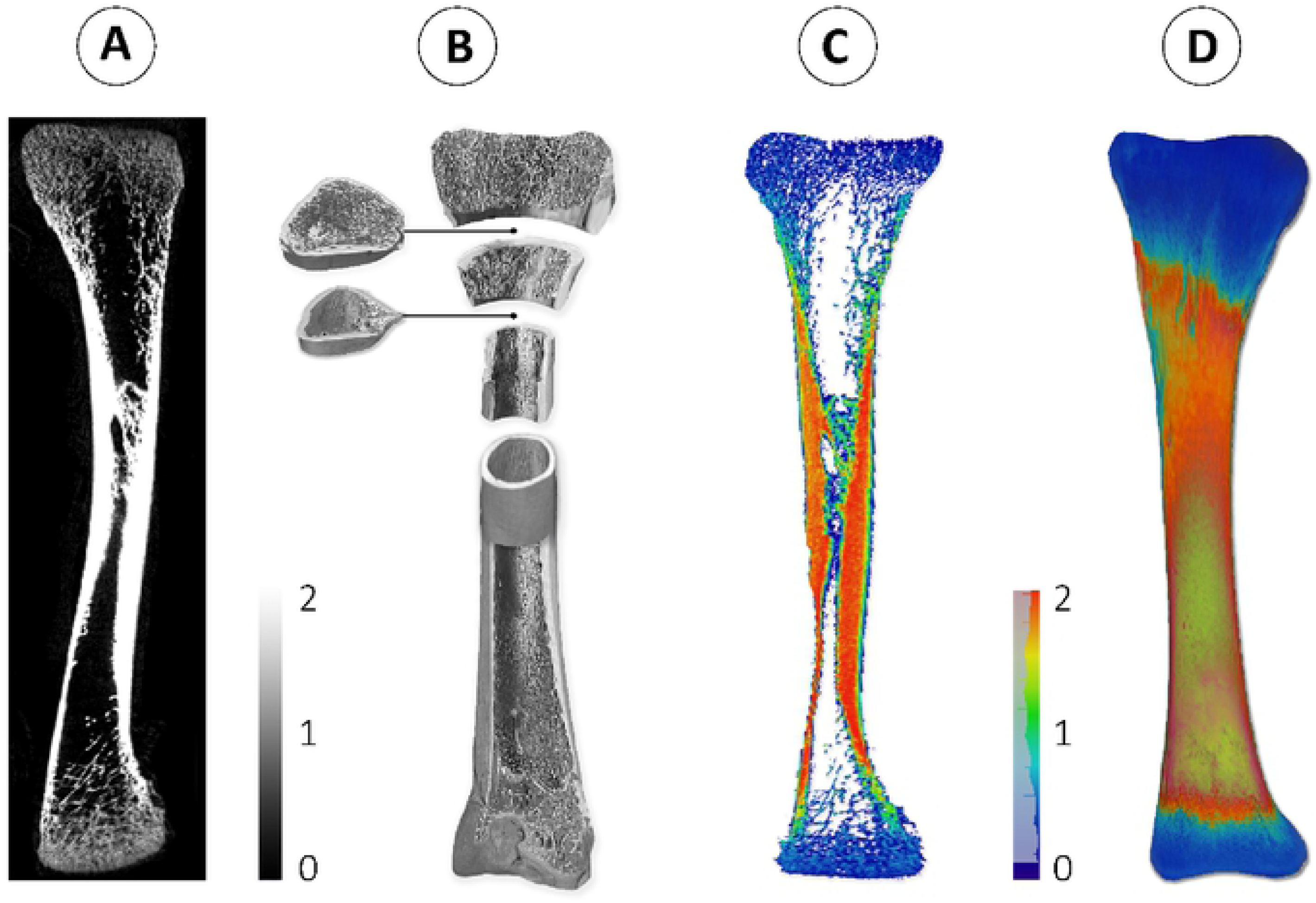
Illustrations of scanned bone by GE Phoenix 3D X-ray microfocus CT scanner visualized in Avizo 3D viewer software. A) Two-dimensional black and white (grey scale) tibia middle slice view. Shades of grey represent the mineralization areas of bone from dark grey (less mineralization) to white (more mineralization). B) Three-dimensional black and white (grey scale) tibia inner and outer view. C) Three-dimensional coloured tibia outer layer view. Colour scale represents the mineralization areas of outer bone from dark blue (less mineralization, 0) to red (more mineralization). D) Three-dimensional coloured tibia middle slice view. Colour scale represents the mineralization areas of bone from blue (less mineralization, 0) to green (more mineralization, 2).

The same tibia bones used for 3D X-ray scanning were subjected to a three-point bending test, of which the method is described by [58], using an Instron^®^ electromechanical universal testing machine (Instron^®^, Norwood, Massachusetts, United States). Maximal load of breaking point was used as the tibia ultimate strength; reached yield load just before the angle has changed on slope data was used as the tibia yield strength; the slope of the selected linear part of the curve data was used as the tibia stiffness; the area under the curve of selected region data was used as the tibia energy to fracture. Elastic modulus (GPa), which is the amount of strain as a result of a particular amount of stress [59], was calculated using the following formula [59, 60]:

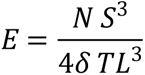

where *E* is the elastic modulus (GPa), *N* is the maximal load (N), S is the span between bending fixtures (mm), T is the tibia thickness (mm), L is the tibia length (mm), and δ is the maximum deflection (mm) at the midpoint of the bone.

### Statistical Analysis

All statistical analyses were performed in SAS (Version 9.4, 2013, SAS Institute Inc., Cary, North Carolina, US). Model assumptions were approved at both means and residuals for continuous data. Non-normal distributed data were log-transformed before analyses. Pen was used as the experimental unit for all analyses.

All growth performance data from day 0 to 35 (BW, FI, FCR, mortality) were subjected to general mixed model analysis, using the MIXED procedure with model 1.

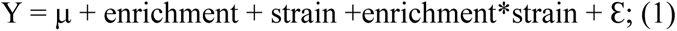

where Y = the dependent variable, μ = the overall mean, enrichment = whether or not pen enrichment was applied (enriched or non-enriched), strain = broiler strain (fast-growing Ross 308 or slower-growing Hubbard JA757), interaction = 2-way interaction between enrichment and strain, Ɛ = residual error.

From day 35 onwards, only chickens from the slower-growing strain were present and growth performance data (BW, FI, FCR, mortality) was subjected to general linear mixed model analysis, using the MIXED procedure with model 2.

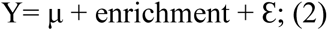

where Y = the dependent variable, μ is the overall mean, enrichment = whether or not pen enrichment was applied (enriched or non-enriched), Ɛ = residual error.

Tibia morphological, biophysical and mechanical characteristics, at two body weight classes (1400 and 2200 g), were subjected to general linear mixed model analysis, using the MIXED procedure with model 1.

Home pen behaviour (eating, drinking, walking, standing, sitting, comfort behaviour, foraging, dustbathing, ground pecking, aggression and others) and gait score were subjected to general linear mixed model analysis, using the MIXED procedure with model 1 (home pen behaviour at day 8, 22 and 29) and model 2 (home pen behaviour at day 43). Preliminary analyses demonstrated a lack of interaction between strain and enrichment for home pen behaviour and consequently data is presented for only main effects. Gait score at day 27 (fast-growing chickens) and day 35 (slower-growing chickens), when they had similar body weights, were compared, using model 1, by including actual BW as a covariate.

Use of enrichment (chickens on platforms and ramps, chickens under platforms and ramps, dustbathing chickens and chickens perching on barriers) was subjected to general linear mixed model analysis, using the MIXED procedure with model 3.

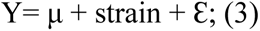

where Y = the dependent variable, μ is the overall mean, strain = broiler breeder strain (fast-growing Ross 308 or slower-growing Hubbard JA757), Ɛ = residual error.

To eliminate BW effect between the fast and slower-growing strain, home pen behaviour and enrichment use at day 22 for fast-growing chickens and day 29 for slower-growing chickens, when they had similar body weights, were compared, using model 1 (home pen behaviour) or model 3 (enrichment).

VV^R^ and VV^L^ were subjected to general linear mixed model analysis, at two body weight classes (1400 and 2200 gram), using the MIXED procedure with model 1. Leg disorders (TD, EPA, BCO and EPI) were subjected to generalized linear mixed model analysis, at two body weight classes (1400 and 2200 gram), using the GLIMMIX procedure with model 1. Leg disorders were scored as 0 (no abnormalities), 1 (minor abnormality), or 2 (severe abnormality), but analyzed as 0 (no abnormalities) or 1 (abnormalities present). EPA, BCO and EPI were not statistically analysed, because there were only three chickens scored with BCO and no observations at all were recorded for EPA and EPI.

Results are provided as LSmeans ± SEM, unless indicated otherwise. When multiple comparisons were performed, the level of significance was corrected, using Bonferroni. Effects were considered to be significant at P≤0.05.

## Results

### Performance

No interaction effects between enrichment and strain were found on BW (Table 1). Chickens in non-enriched pens had a higher BW than chickens in enriched pens at day 21 (Δ=35 g, *P*=0.02), 28 (Δ=62 g, *P*=0.007), 35 (Δ=99 g, *P*= 0.003), 42 (slower-growing chickens only; Δ=84 g, *P*=0.003) and 49 (slower-growing chickens only; Δ=93 g, *P*=0.005). Slower-growing broilers had a lower BW than fast-growing broilers at day 0 (Δ=1.8 g), 7 (Δ=29 g), 14 (Δ=134 g), 21 (Δ=321 g), 28 (Δ=540 g) and 35 (Δ=822 g) (all *P*<0.001).

**Table 1.** Effects of pen enrichment (enriched or non-enriched), broiler strain (fast-growing Ross 308 or slower-growing Hubbard JA 757) and their interaction on body weight (gram) of male broiler chickens at different ages (n=7 pens per treatment, LSmeans±SEM).

| Parameter | Day 0 | Day 7 | Day 14 | Day 21 | Day 28 | Day 35 | Day 42<sup>1</sup> | Day 49<sup>1</sup> |
| --- | --- | --- | --- | --- | --- | --- | --- | --- |
| Enrichment |  |  |  |  |  |  |  |  |
| Enriched | 36.9 | 124 | 334 | 657<sup>b</sup> | 1070<sup>b</sup> | 1596<sup>b</sup> | - | - |
| Non-enriched | 36.9 | 126 | 346 | 692<sup>a</sup> | 1132<sup>a</sup> | 1695<sup>a</sup> | - | - |
| SEM | 0.1 | 3 | 6 | 10 | 15 | 20 | - | - |
| Strain |  |  |  |  |  |  |  |  |
| Fast | 37.8<sup>a</sup> | 140<sup>a</sup> | 407<sup>a</sup> | 835<sup>a</sup> | 1371<sup>a</sup> | 2057<sup>a</sup> | - | - |
| Slower | 36.0<sup>b</sup> | 111<sup>b</sup> | 273<sup>b</sup> | 514<sup>b</sup> | 831<sup>b</sup> | 1235<sup>b</sup> | - | - |
| SEM | 0.1 | 3 | 6 | 10 | 15 | 20 | - | - |
| Enrichment*strain |  |  |  |  |  |  |  |  |
| Enriched fast | 37.8 | 140 | 401 | 817 | 1324 | 1985 | - | - |
| Non-enriched fast | 37.8 | 140 | 413 | 854 | 1419 | 2129 | - | - |
| Enriched slower | 36.0 | 108 | 267 | 497 | 817 | 1208 | 1641<sup>b</sup> | 2144<sup>b</sup> |
| Non-enriched slower | 36.0 | 113 | 279 | 530 | 845 | 1261 | 1724<sup>a</sup> | 2237<sup>a</sup> |
| SEM | 0.2 | 4 | 8 | 14 | 21 | 29 | 15 | 19 |
| *P*-values |  |  |  |  |  |  |  |  |
| Enrichment | 0.78 | 0.60 | 0.17 | 0.02 | 0.007 | 0.003 | 0.003 | 0.005 |
| Strain | <0.001 | <0.001 | <0.001 | <0.001 | <0.001 | <0.001 | - | - |
| Enrichment*strain | 0.88 | 0.55 | 0.99 | 0.88 | 0.13 | 0.13 | - | - |

<sup>a-b</sup>Values within a column and factor lacking a common superscript differ ( $P \leq 0.05$ ).
<sup>1</sup>At day 42 and 49, only chickens of the slower-growing strain were present.

No interaction effects between pen enrichment and strain were found on FI (Table 2) and neither pen enrichment effects were found. Slower-growing chickens had a lower FI than fast-growing broilers between day 0-14 (Δ=112 g), day 14-35 (Δ=923 g) and day 0-35 (Δ=1034 g) (all *P*<0.001).

**Table 2.** Effects of pen enrichment (enriched or non-enriched), broiler strain (fast-growing Ross 308 or slower-growing Hubbard JA 757) and their interaction on feed intake (FI; gram per chicken) and feed conversion ratio (FCR=FI/BWG^1^) of male broiler chickens in different phases of the rearing period (n=7 pens per treatment, LSmeans±SEM).

| Parameter | FI<br>d<br>0-14 | FI<br>d<br>14-35 | FI<br>d<br>0-35 | FI<br>d<br>35-49<sup>2</sup> | FI<br>d<br>0-49<sup>2</sup> | FCR<br>d<br>0-14 | FCR<br>d<br>14-35 | FCR<br>d<br>0-35 | FCR<br>d<br>35-49<sup>2</sup> | FCR<br>d<br>0-49<sup>2</sup> |
| --- | --- | --- | --- | --- | --- | --- | --- | --- | --- | --- |
| Enrichment |  |  |  |  |  |  |  |  |  |  |
| Enriched | 350 | 2050 | 2400 | - | - | 1.20<sup>a</sup> | 1.62<sup>a</sup> | 1.54<sup>a</sup> | - | - |
| Non-enriched | 344 | 2110 | 2470 | - | - | 1.13<sup>b</sup> | 1.57<sup>b</sup> | 1.49<sup>b</sup> | - | - |
| SEM | 7 | 22 | 27 | - | - | 0.01 | 0.01 | 0.01 | - | - |
| Strain |  |  |  |  |  |  |  |  |  |  |
| Fast | 403<sup>a</sup> | 2528<sup>a</sup> | 2930<sup>b</sup> | - | - | 1.10<sup>b</sup> | 1.53<sup>b</sup> | 1.45<sup>b</sup> | - | - |
| Slower | 291<sup>b</sup> | 1605<sup>b</sup> | 1896<sup>a</sup> | - | - | 1.23<sup>a</sup> | 1.67<sup>a</sup> | 1.58<sup>a</sup> | - | - |
| SEM | 7 | 22 | 27 | - | - | 0.01 | 0.01 | 0.01 | - | - |
| Enrichment*strain |  |  |  |  |  |  |  |  |  |  |
| Enriched fast | 408 | 2480 | 2882 | - | - | 1.13 | 1.56 | 1.48 | - | - |
| Non-enriched fast | 398 | 2574 | 2971 | - | - | 1.07 | 1.50 | 1.42 | - | - |
| Enriched slower | 291 | 1589 | 1895 | 2029 | 3909 | 1.26 | 1.69 | 1.61 | 2.17 | 1.85<sup>a</sup> |
| Non-enriched slower | 291 | 1620 | 1916 | 2037 | 3948 | 1.20 | 1.65 | 1.56 | 2.09 | 1.79<sup>b</sup> |
| SEM | 10 | 31 | 38 | 14 | 23 | 0.01 | 0.01 | 0.01 | 0.02 | 0.01 |
| P-values |  |  |  |  |  |  |  |  |  |  |
| Enrichment | 0.60 | 0.15 | 0.29 | 0.72 | 0.26 | <0.001 | <0.001 | <0.001 | 0.06 | <0.001 |
| Strain | <0.001 | <0.001 | <0.001 | - | - | <0.001 | <0.001 | <0.001 | - | - |
| Enrichment*strain | 0.58 | 0.63 | 0.81 | - | - | 0.74 | 0.47 | 0.53 | - | - |

<sup>a-b</sup>Values within a column and factor lacking a common superscript differ (P≤0.05).
<sup>1</sup>Body weight gain.
<sup>2</sup>Only slower-growing chickens.
\*FI was calculated without BSFL intake of chickens in enriched pens and by excluding the intake of the dietary supplement of chickens in non-enriched pens.

No interaction effects between enrichment and strain were found on FCR (Table 2). Chickens in non-enriched pens had a lower FCR than chickens in enriched pens between days 0-14 (Δ=0.07), days 14-35 (Δ=0.05), days 0-35 (Δ=0.05) and days 0-49 (Δ=0.05) (all *P*<0.001). Slower-growing chickens had a higher FCR than fast-growing chickens between days 0-14 (Δ=0.13), 14-35 (Δ=0.14) and 0-35 (Δ=0.13) (all *P*<0.001).

### Tibia morphological characteristics

At the 1400 gram BW class, no interaction effects between pen enrichment and strain were found on tibia morphological characteristics (Table 3) and neither pen enrichment effects were found. Slower-growing chickens had a higher femoral (Δ=0.17 cm, *P*=0.02) and metatarsal side proximal tibia head thicknesses (Δ=0.12 cm, *P*=0.04) than fast-growing chickens. At the 2200 gram BW class, no interaction effects between enrichment and strain were found on tibia morphological characteristics (Table 3) and neither pen enrichment effects were found. Slower-growing broilers had a higher tibia weight (Δ=0.81 g, *P*=0.02), proximal tibia length (Δ=0.63 cm, *P*=0.008) and metatarsal side proximal tibia head thicknesses (Δ=0.17 cm, *P*=0.002) than fast-growing broilers.

**Table 3.**
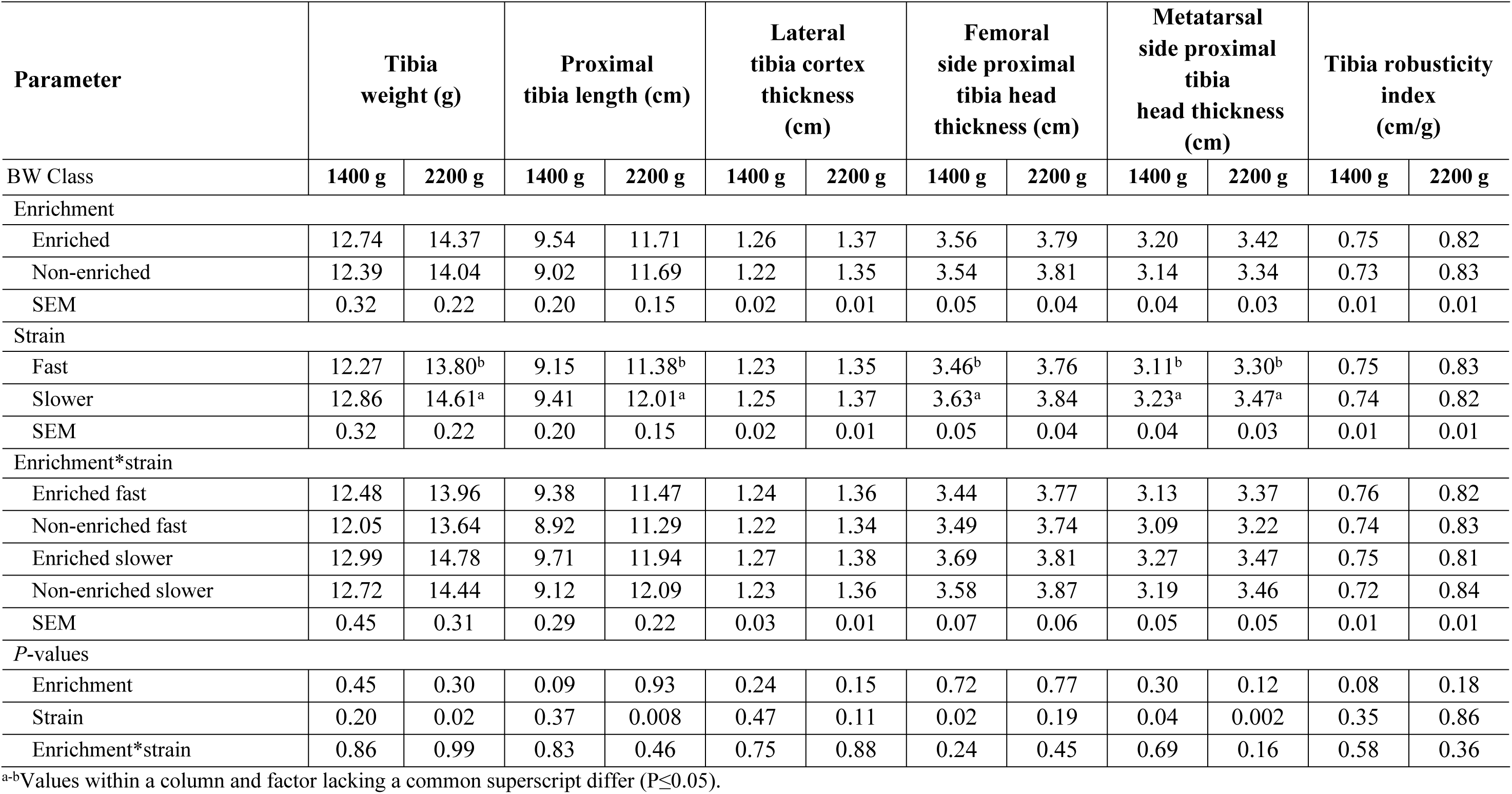
Effects of pen enrichment (enriched or non-enriched), broiler strain (fast-growing Ross 308 or slower-growing Hubbard JA 757) and their interaction on tibia morphological characteristics of male broiler chickens in two body weight classes (1400 and 2200 gram) (2 chickens per pen, n=7 pens per treatment, LSmeans±SEM).

### Tibia biophysical characteristics

At the 1400 gram BW class, no interaction effects between pen enrichment and strain were found on tibia biophysical characteristics (Table 4) and neither pen enrichment effects were found. Slower-growing broilers had a higher tibia osseous volume (Δ=6.4 cm^3^, *P*<0.001), tibia total volume (Δ=6.6 cm^3^, *P*<0.001), tibia volume fraction (Δ=0.04 %, *P*<0.001) and tibia mineral content (Δ=1.1 g, *P*<0.001) than fast-growing broilers. At the 2200 gram BW class, an interaction between pen enrichment and strain was found on tibia pore volume (Table 4). Enriched slower-growing group resulted in a lower tibia pore volume compared to other three groups (Δ=1.0 cm^3^ on average; *P*=0.02). Chickens in non-enriched pens had a lower tibia osseous volume (Δ=1.8 cm^3^, *P*=0.003), tibia total volume (Δ=1.4 cm^3^, *P*=0.03) and tibia volume fraction (Δ=0.02 %, *P*=0.002) than chickens in enriched pens. Slower-growing broilers had a higher tibia osseous volume (Δ=5.9 cm^3^, *P*<0.001), tibia total volume (Δ=5.4 cm^3^, *P*<0.001), tibia volume fraction (Δ=0.05 %, *P*<0.001), tibia mineral content (Δ=0.7 g, *P*=0.02) and tibia mineral density (Δ=0.05 g/cm^3^, *P*<0.001) than fast-growing broilers.

**Table 4.** Effects of pen enrichment (enriched or non-enriched), broiler strain (fast-growing Ross 308 or slower-growing Hubbard JA 757) and their interaction on tibia biophysical characteristics of male broiler chickens in two body weight classes (1400 and 2200 gram) (2 chickens per pen, n=7 pens per treatment, LSmeans±SEM).

| Parameter | Tibia osseous volume (cm<sup>3</sup>) |  | Tibia pore volume (cm<sup>3</sup>) |  | Tibia total volume (cm<sup>3</sup>) |  | Tibia volume fraction (OV<sup>1</sup>/TV<sup>2</sup>) |  | Tibia mineral content (g) |  | Tibia mineral density (g/cm<sup>3</sup>) |  |
| --- | --- | --- | --- | --- | --- | --- | --- | --- | --- | --- | --- | --- |
|  | 1400 g | 2200 g | 1400 g | 2200 g | 1400 g | 2200 g | 1400 g | 2200 g | 1400 g | 2200 g | 1400 g | 2200 g |
| BW Class |  |  |  |  |  |  |  |  |  |  |  |  |
| Enrichment |  |  |  |  |  |  |  |  |  |  |  |  |
| Enriched | 21.3 | 25.3<sup>a</sup> | 3.5 | 3.9 | 24.8 | 29.2<sup>a</sup> | 0.86 | 0.86<sup>a</sup> | 11.2 | 12.5 | 0.21 | 0.37 |
| Non-enriched | 20.7 | 23.5<sup>b</sup> | 3.8 | 4.3 | 24.5 | 27.8<sup>b</sup> | 0.84 | 0.84<sup>b</sup> | 11.3 | 12.0 | 0.19 | 0.37 |
| SEM | 0.6 | 0.4 | 0.2 | 0.1 | 0.6 | 0.4 | 0.01 | 0.01 | 0.2 | 0.2 | 0.01 | 0.01 |
| Strain |  |  |  |  |  |  |  |  |  |  |  |  |
| Fast | 17.8<sup>b</sup> | 21.4<sup>a</sup> | 3.5 | 4.4 | 21.3<sup>b</sup> | 25.8<sup>b</sup> | 0.83<sup>b</sup> | 0.83<sup>b</sup> | 10.7<sup>b</sup> | 11.9<sup>b</sup> | 0.19 | 0.34<sup>b</sup> |
| Slower | 24.2<sup>a</sup> | 27.3<sup>b</sup> | 3.7 | 3.9 | 27.9<sup>a</sup> | 31.2<sup>a</sup> | 0.87<sup>a</sup> | 0.88<sup>a</sup> | 11.8<sup>a</sup> | 12.6<sup>a</sup> | 0.21 | 0.39<sup>a</sup> |
| SEM | 0.6 | 0.4 | 0.2 | 0.1 | 0.6 | 0.4 | 0.01 | 0.01 | 0.2 | 0.2 | 0.01 | 0.01 |
| Enrichment*strain |  |  |  |  |  |  |  |  |  |  |  |  |
| Enriched fast | 18.6 | 22.5 | 3.5 | 4.4<sup>a</sup> | 22.1 | 26.9 | 0.84 | 0.84 | 10.5 | 12.0 | 0.20 | 0.35 |
| Non-enriched fast | 17.0 | 20.3 | 3.6 | 4.4<sup>a</sup> | 20.6 | 24.7 | 0.83 | 0.82 | 10.9 | 11.9 | 0.18 | 0.34 |
| Enriched slower | 24.0 | 28.0 | 3.5 | 3.4<sup>b</sup> | 27.4 | 31.5 | 0.87 | 0.89 | 11.9 | 12.9 | 0.22 | 0.39 |
| Non-enriched slower | 24.5 | 26.6 | 3.9 | 4.3<sup>a</sup> | 28.4 | 30.9 | 0.86 | 0.86 | 11.7 | 12.2 | 0.21 | 0.40 |
| SEM | 0.8 | 0.5 | 0.2 | 0.2 | 0.9 | 0.6 | 0.01 | 0.01 | 0.2 | 0.2 | 0.01 | 0.01 |
| P-values |  |  |  |  |  |  |  |  |  |  |  |  |
| Enrichment | 0.51 | 0.003 | 0.18 | 0.02 | 0.79 | 0.03 | 0.10 | 0.002 | 0.65 | 0.07 | 0.21 | 0.78 |
| Strain | <0.001 | <0.001 | 0.50 | 0.008 | <0.001 | <0.001 | <0.001 | <0.001 | <0.001 | 0.02 | 0.08 | <0.001 |
| Enrichment*strain | 0.20 | 0.50 | 0.51 | 0.02 | 0.19 | 0.16 | 0.74 | 0.17 | 0.31 | 0.26 | 0.59 | 0.19 |

<sup>a-b</sup>Values within a column and factor lacking a common superscript differ (P≤0.05).
<sup>1</sup>Osseous volume.
<sup>2</sup>Total volume.

### Tibia mechanical characteristics

At the 1400 gram BW class, no interaction effects between pen enrichment and strain were found on tibia mechanical characteristics and neither pen enrichment effects were found (Table 5). Slower-growing broilers had a higher tibia ultimate strength (Δ=21.7 N, *P*<0.001), tibia yield strength (Δ=21.0 N, *P*<0.001), tibia stiffness (Δ=20.6 N/mm, *P*<0.001) and tibia energy to fracture (Δ=21.9 N-mm, *P*<0.001) than fast-growing broilers. At the 2200 gram BW class, no interaction effects between pen enrichment and strain were found on tibia mechanical characteristics (Table 5) and neither pen enrichment effects were found. Slower-growing chickens had a higher tibia ultimate strength (Δ=19.4 N, *P*<0.001), tibia yield strength (Δ=17.8 N, *P*<0.001), tibia stiffness (Δ=21.7 N/mm, *P*<0.001) and tibia energy to fracture (Δ=20.9 N-mm, *P*<0.001) than fast-growing broilers.

**Table 5.** Effects of pen enrichment (enriched or non-enriched), broiler breeder strain (fast-growing Ross 308 or slower-growing Hubbard JA 757), and their interaction on tibia mechanical characteristics of male broiler offspring in two body weight classes (1400 and 2200 gram) (2 chickens per pen, n=7 pens per treatment, LSmeans±SEM).

| Parameter | Tibia ultimate strength (N) |  | Tibia yield strength (N) |  | Tibia stiffness (N/mm) |  | Tibia energy to fracture (N-mm) |  | Tibia elastic modulus (GPa) |  |
| --- | --- | --- | --- | --- | --- | --- | --- | --- | --- | --- |
|  | 1400 g | 2200 g | 1400 g | 2200 g | 1400 g | 2200 g | 1400 g | 2200 g | 1400 g | 2200 g |
| BW Class |  |  |  |  |  |  |  |  |  |  |
| Enrichment |  |  |  |  |  |  |  |  |  |  |
| Enriched | 242.4 | 274.6 | 226.4 | 256.1 | 233.5 | 269.1 | 231.4 | 261.9 | 12.6 | 12.2 |
| Non-enriched | 237.2 | 271.2 | 223.4 | 254.3 | 228.6 | 263.8 | 226.8 | 260.2 | 12.4 | 12.0 |
| SEM | 2.2 | 1.8 | 2.1 | 2.1 | 2.1 | 1.9 | 2.0 | 1.9 | 0.4 | 0.3 |
| Strain |  |  |  |  |  |  |  |  |  |  |
| Fast | 229.0<sup>b</sup> | 263.2<sup>b</sup> | 214.4<sup>b</sup> | 246.3<sup>b</sup> | 220.7<sup>b</sup> | 255.6<sup>b</sup> | 218.1<sup>b</sup> | 250.6<sup>b</sup> | 12.4 | 12.6 |
| Slower | 250.6<sup>a</sup> | 282.6<sup>a</sup> | 235.4<sup>a</sup> | 264.1<sup>a</sup> | 241.3<sup>a</sup> | 277.3<sup>a</sup> | 240.0<sup>a</sup> | 271.5<sup>a</sup> | 12.6 | 11.6 |
| SEM | 2.2 | 1.8 | 2.1 | 2.1 | 2.1 | 1.9 | 2.0 | 1.9 | 0.4 | 0.3 |
| Enrichment*strain |  |  |  |  |  |  |  |  |  |  |
| Enriched fast | 229.0 | 263.4 | 213.5 | 246.2 | 220.6 | 258.9 | 217.9 | 250.5 | 12.4 | 13.0 |
| Non-enriched fast | 229.1 | 263.0 | 215.4 | 246.4 | 220.9 | 252.3 | 218.3 | 250.7 | 12.4 | 12.1 |
| Enriched slower | 255.9 | 285.7 | 239.3 | 266.0 | 246.4 | 279.3 | 244.8 | 273.3 | 12.7 | 11.5 |
| Non-enriched slower | 245.4 | 279.4 | 231.5 | 262.1 | 236.3 | 275.3 | 235.3 | 269.8 | 12.5 | 11.8 |
| SEM | 3.1 | 2.6 | 2.9 | 3.0 | 2.9 | 2.7 | 2.9 | 2.7 | 0.6 | 0.4 |
| P-values |  |  |  |  |  |  |  |  |  |  |
| Enrichment | 0.11 | 0.22 | 0.32 | 0.55 | 0.11 | 0.06 | 0.13 | 0.55 | 0.83 | 0.55 |
| Strain | <0.001 | <0.001 | <0.001 | <0.001 | <0.001 | <0.001 | <0.001 | <0.001 | 0.78 | 0.06 |
| Enrichment*strain | 0.10 | 0.27 | 0.11 | 0.49 | 0.09 | 0.64 | 0.10 | 0.49 | 0.90 | 0.18 |

<sup>a-b</sup>Values within a column and factor lacking a common superscript differ ( $P \leq 0.05$ ).

### Leg disorders and gait score

No interaction effects between pen enrichment and broiler breeder strain were found on VV^R^, VV^L^ and TD at both 1400 and 2200 gram BW classes and no main effects of pen enrichment or strain were found on TD. At the 1400 gram BW class, VV angulation was not affected by enrichment or strain (on average 4.90°). At the 2200 gram BW class, slower-growing chickens had a lower VV^R^ than fast-growing chickens (3.80° vs 6.04°, respectively, *P*=0.003). VV^L^ angulation was not affected by enrichment or strain at this BW class (on average 5.88°). No interaction and main effects were found on TD, which was observed in 11 chickens (9.82 %) at 1400 gram BW class and 10 chickens (8.92 %) at 2200 BW class.

At similar BW class (day 27 for fast-growing chickens and day 35 for slower-growing chickens), an interaction between pen enrichment and strain was found on gait score. The NS group had a lower gait score compared to NF group (1.66 vs. 2.11, respectively; *P*=0.003), while other two groups were in between (both 1.93).

### Home pen behaviour and use of enrichment

Chickens in non-enriched pens showed less foraging behaviour at day 8 (Δ=8.47 %, *P*<0.001), day 22 (Δ=9.19 %, *P*<0.001), day 29 (Δ=5.6 %, *P*<0.001) and day 43 (slower-growing chickens only, Δ=8.86 %, *P*<0.001), less dust bathing behaviour at day 8 (Δ=0.97 %, *P*=0.006) and less ground pecking behaviour at day 43 (slower-growing chickens only, Δ=3.98 %, *P*=0.02) than chickens in enriched pens. The opposite was found for standing behaviour at day 29 (Δ=4.15 %, *P*=0.007), sitting behaviour at day 22 (Δ=8.97 %, *P*=0.03), ground pecking behaviour at day 8 (Δ=3.71 %, *P*=0.002) and aggression behaviour at day 29 (Δ=1.05 %, *P*=0.02), which behaviours were all higher in non-enriched pens than in enriched pens. Slower-growing chickens showed more walking behaviour at day 8 (Δ=6.99 %, *P*=0.001), day 22 (Δ=7.6 %, *P*<0.001) and day 29 (Δ=5.32 %, *P*<0.001), more standing behaviour at day 8 (Δ=2.34 %, *P*=0.03), day 22 (Δ=3.12 %, *P*=0.02) and day 29 (Δ=8.91 %, *P*<0.001), more foraging behaviour at day 22 (Δ=6.61 %, *P*=0.009) and day 29 (Δ=3.9 %, *P*=0.002) and more aggression behaviour at day 22 (Δ=0.94 %, *P*=0.03) and day 29 (Δ=1.02 %, *P*=0.03) than fast-growing chickens. The opposite was found for eating behaviour at day 8 (Δ=3.04 %, *P*=0.04) and day 22 (Δ=1.16 %, *P*=0.03) and sitting behaviour at day 22 (Δ=19.28 %, *P*<0.001) and day 29 (Δ=8.91 %, *P*<0.001).

In enriched pens, a clear strain effect was found on use of different enrichment objects (Table 7). The percentage of chickens on platforms and ramps at day 29 (Δ=14.6 %, *P*<0.001) and perching on barriers at day 8 (Δ=4.9 %, *P*<0.001), day 22 (Δ=14.05 %, *P*<0.001) and day 29 (Δ=16.05 %, *P*<0.001) was higher in slower-growing chickens than in fast-growing chickens. The opposite was found for the percentage chickens under platforms and ramps at day 29 (Δ=13.72 %, *P*=0.003) and dustbathing chickens at day 8 (Δ=1.19 %, *P*=0.05).

**Table 6.** Effects of pen enrichment (enriched or non-enriched) and broiler strain (fast-growing Ross 308 or slower-growing Hubbard JA 757) on percentage of male chickens showing eating, drinking, walking, standing, sitting, comfort behaviour, foraging, dustbathing, ground pecking, aggression and other behaviours at day 8, 22, 29 and 43 of age (n=7 pens per treatment; LSmeans±SEM).

| Parameter and day (%) | Pen enrichment |  | Strain |  | SEM | *P*-values<sup>2</sup> |  |
| --- | --- | --- | --- | --- | --- | --- | --- |
|  | Enriched | Non-enriched | Fast | Slower |  | Enrichment | Strain |
| Eating |  |  |  |  |  |  |  |
| Day 8 | 6.90 | 6.06 | 8.00<sup>b</sup> | 4.96<sup>a</sup> | 0.94 | 0.54 | 0.04 |
| Day 22 | 1.65 | 1.80 | 2.31<sup>b</sup> | 1.15<sup>a</sup> | 0.33 | 0.75 | 0.03 |
| Day 29 | 1.78 | 2.78 | 2.86 | 1.69 | 0.46 | 0.14 | 0.09 |
| Day 43<sup>1</sup> | 1.65 | 1.79 | - | - | 0.51 | 0.86 | - |
| Drinking |  |  |  |  |  |  |  |
| Day 8 | 13.17 | 12.01 | 13.76 | 11.42 | 1.58 | 0.61 | 0.31 |
| Day 22 | 5.33 | 5.58 | 5.67 | 5.24 | 0.72 | 0.81 | 0.69 |
| Day 29 | 4.36 | 4.78 | 5.00 | 4.13 | 0.72 | 0.69 | 0.40 |
| Day 43<sup>1</sup> | 3.78 | 3.23 | - | - | 0.58 | 0.51 | - |
| Walking |  |  |  |  |  |  |  |
| Day 8 | 13.42 | 12.27 | 9.35<sup>b</sup> | 16.34<sup>a</sup> | 1.10 | 0.47 | 0.001 |
| Day 22 | 7.92 | 7.09 | 3.70<sup>b</sup> | 11.30<sup>a</sup> | 1.22 | 0.64 | <0.001 |
| Day 29 | 5.67 | 6.90 | 3.62<sup>b</sup> | 8.94<sup>a</sup> | 0.82 | 0.31 | <0.001 |
| Day 43<sup>1</sup> | 7.99 | 8.56 | - | - | 1.46 | 0.80 | - |
| Standing |  |  |  |  |  |  |  |
| Day 8 | 5.68 | 7.65 | 5.50<sup>b</sup> | 7.84<sup>a</sup> | 0.72 | 0.07 | 0.03 |
| Day 22 | 4.54 | 5.44 | 3.43<sup>b</sup> | 6.55<sup>a</sup> | 0.81 | 0.44 | 0.02 |
| Day 29 | 5.50<sup>b</sup> | 9.65<sup>a</sup> | 3.12<sup>b</sup> | 12.03<sup>a</sup> | 1.00 | 0.007 | <0.001 |
| Day 43<sup>1</sup> | 13.64 | 15.84 | - | - | 1.46 | 0.29 | - |
| Sitting |  |  |  |  |  |  |  |
| Day 8 | 32.57 | 37.36 | 37.61 | 33.31 | 2.44 | 0.18 | 0.14 |
| Day 22 | 45.61<sup>b</sup> | 54.58<sup>a</sup> | 59.73<sup>a</sup> | 40.45<sup>b</sup> | 2.72 | 0.03 | <0.001 |
| Day 29 | 57.36 | 55.23 | 64.78<sup>a</sup> | 47.80<sup>b</sup> | 2.90 | 0.61 | <0.001 |
| Day 43<sup>1</sup> | 50.06 | 58.22 | - | - | 3.97 | 0.18 | - |
| Comfort behaviour |  |  |  |  |  |  |  |
| Day 8 | 7.55 | 9.34 | 7.15 | 9.74 | 1.00 | 0.22 | 0.08 |
| Day 22 | 8.86 | 10.12 | 8.84 | 10.13 | 1.08 | 0.42 | 0.41 |
| Day 29 | 6.19 | 7.60 | 7.27 | 6.52 | 1.12 | 0.39 | 0.64 |
| Day 43<sup>1</sup> | 4.46 | 3.65 | - | - | 0.95 | 0.56 | - |
| Foraging |  |  |  |  |  |  |  |
| Day 8 | 14.45<sup>a</sup> | 5.98<sup>b</sup> | 9.67 | 10.76 | 1.00 | <0.001 | 0.45 |
| Day 22 | 12.88<sup>a</sup> | 3.69<sup>b</sup> | 4.98<sup>b</sup> | 11.59<sup>a</sup> | 1.64 | <0.001 | 0.009 |
| Day 29 | 8.50<sup>a</sup> | 2.90<sup>b</sup> | 3.75<sup>b</sup> | 7.65<sup>a</sup> | 0.79 | <0.001 | 0.002 |
| Day 43<sup>1</sup> | 9.78<sup>a</sup> | 0.92<sup>b</sup> | - | - | 1.25 | <0.001 | - |
| Dust bathing |  |  |  |  |  |  |  |
| Day 8 | 1.09<sup>a</sup> | 0.12<sup>b</sup> | 0.90 | 0.30 | 0.23 | 0.006 | 0.07 |
| Day 22 | 0.65 | 0.44 | 0.65 | 0.44 | 0.24 | 0.56 | 0.54 |
| Day 29 | 0.46 | 0.67 | 0.36 | 0.77 | 0.28 | 0.60 | 0.31 |
| Day 43<sup>1</sup> | 0.28 | 0.15 | - | - | 0.17 | 0.59 | - |
| Ground pecking |  |  |  |  |  |  |  |
| Day 8 | 4.83<sup>b</sup> | 8.54<sup>a</sup> | 7.40 | 5.97 | 0.73 | 0.002 | 0.18 |
| Day 22 | 12.02 | 10.76 | 10.60 | 12.17 | 1.17 | 0.46 | 0.36 |
| Day 29 | 10.08 | 8.33 | 9.09 | 9.31 | 1.17 | 0.31 | 0.90 |
| Day 43<sup>1</sup> | 8.10<sup>a</sup> | 4.12<sup>b</sup> | - | - | 0.95 | 0.02 | - |
| Aggression |  |  |  |  |  |  |  |
| Day 8 | 0.35 | 0.22 | 0.22 | 0.35 | 0.17 | 0.60 | 0.60 |
| Day 22 | 0.61 | 0.59 | 0.14<sup>b</sup> | 1.06<sup>a</sup> | 0.29 | 0.97 | 0.04 |
| Day 29 | 0.12<sup>b</sup> | 1.17<sup>a</sup> | 0.14<sup>b</sup> | 1.16<sup>a</sup> | 0.29 | 0.02 | 0.03 |
| Day 43<sup>1</sup> | 0.26<sup>b</sup> | 3.53<sup>a</sup> | - | - | 0.98 | 0.04 | - |
| Others<sup>3</sup> |  |  |  |  |  |  |  |
| Day 8 | - | 0.13 | 0.13 | - | 0.08 | 0.26 | 0.26 |
| Day 22 | - | - | - | - | - | - | - |
| Day 29 | - | - | - | - | - | - | - |
| Day 43<sup>1</sup> | - | - | - | - | - | - | - |

<sup>a-b</sup>Values within a factor and row lacking a common superscript differ ( $P \leq 0.05$ ).
<sup>1</sup>Only slower-growing chickens.
<sup>2</sup>No interactions between broiler breeder strain and pen enrichment were observed for any of the behaviours at any of the sampling days.
<sup>3</sup>Chickens demonstrating a behaviour other than eating, drinking, walking, standing, sitting, comfort behaviour, foraging, dustbathing, ground pecking and aggression.

**Table 7.**
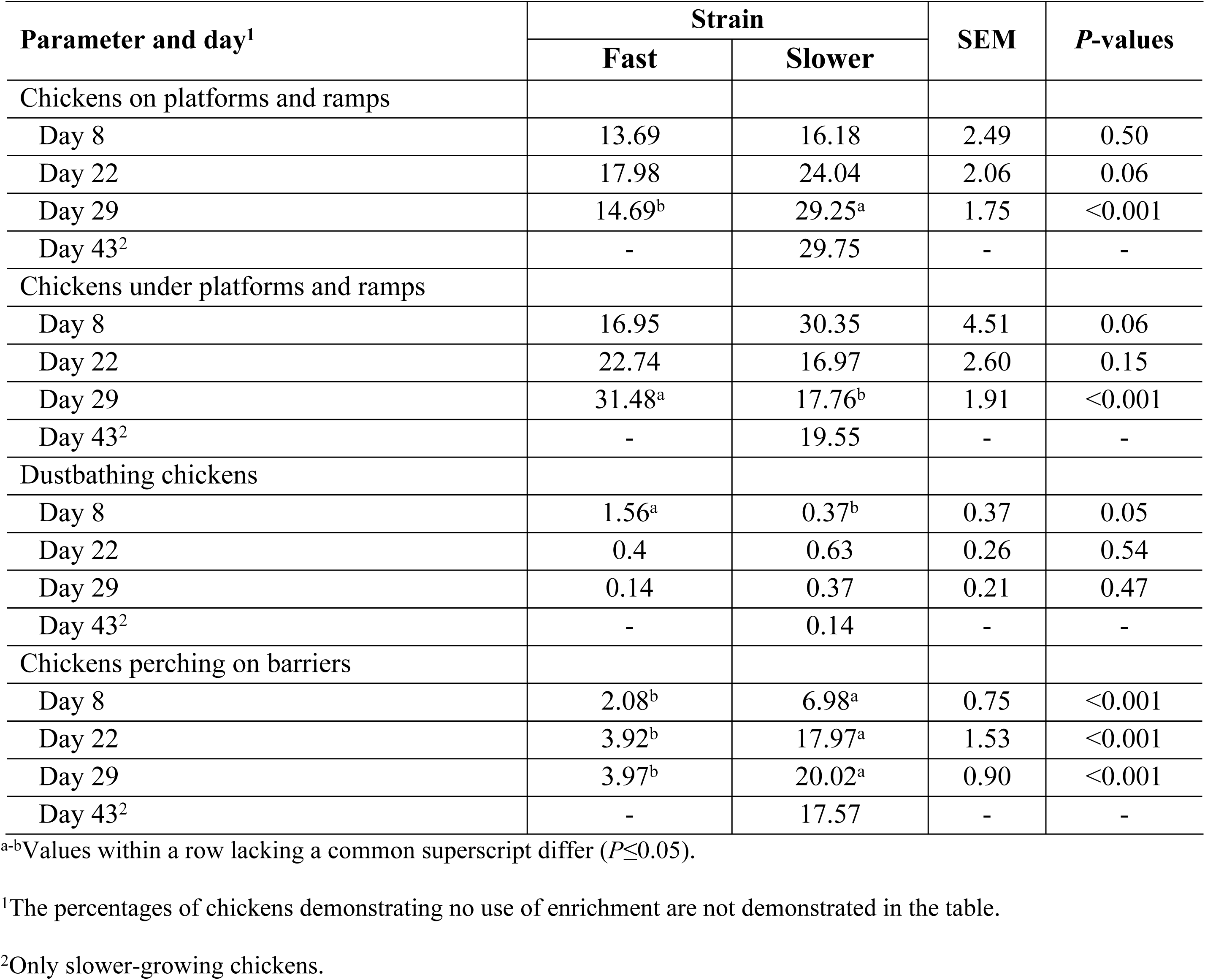
Effects of broiler strain (fast-growing Ross 308 or slower-growing Hubbard JA 757) on percentage of male chickens in enriched pens, using the following enrichment objects (on platforms and ramps, under platforms and ramps, dustbathing, perching on barriers) at day 8, 22, 29 and 43 of age (n=7 pens per treatment, LSmeans±SEM)

At similar BW, fast-growing chickens showed more walking (Δ=5.24 %), standing (Δ=8.6 %), foraging (Δ=2.67 %), dust bathing (Δ=0.12 %) and aggression (Δ=1.02 %) behaviour than fast-growing chickens, while the opposite was found for eating (Δ=0.62 %), drinking (Δ=1.54 %), sitting (Δ=11.93 %) and comfort (Δ=2.32 %) behaviour. At similar BW, more slower-growing chickens were on platforms and ramps (Δ=11.27 %) and perching on barriers (Δ=16.1 %) than fast-growing chickens, whereas the opposite was found for percentage of chickens under platforms and ramps (Δ=4.98 %) and dustbathing chickens (Δ=0.03 %).

## Discussion

The aim of this study was to investigate effects of a combination of different forms of pen enrichment on tibia characteristics, locomotion, leg health and home pen behaviour of both fast and slower-growing broiler chickens. Hardly any interactions were found between strain and enrichment, indicating that both fast and slower-growing chickens are both able to use several forms of enrichment in a comparable way.

### Growth performance

Results of the current study showed that pen enrichment resulted in lower body weight and higher FCR in both fast and slower-growing chickens. These findings are supported by recent studies [61, 62], who observed a negative effect of pen enrichment on growth performance. In an earlier comparable study [38], pen enrichment resulted in lower body weight, but also in a higher FI and a lack of effect on FCR. In the latter study, the FI of the non-enriched pens included the protein-fat mixture, where this was excluded in the current study. Results of current study are not in accordance with other studies [34,63,64,65,66], who found no significant effects of pen enrichment on any growth performance parameters. This discrepancy among studies might be related to the fact that less complex pen enrichment forms were used in these studies compared to the current study. For example, only barrier perches [34,63,64] and mirror, ball, perch and dust (each material in another pen) [65] were used in these studies. The lower body weight gain of enriched-housed broilers in the current experiment might be related to 1) the comprehensive enrichment design of the current study, which contains a combination of platforms, angular ramps, barrier perches, large distance between feed and water and live Black Soldier fly larvae in the moss-peat dust bathing area. A plethora of different enrichments might exponentially stimulate physical activity and consequently a higher metabolic energy use, which will result in a lower body weight gain. This higher activity in enriched pens is supported by the higher percentage of chickens showing active behaviours and use of enrichment in both fast and slower-growing chickens. 2) Chickens might have had difficulties to cross the barrier perches to access the feed on one side and water on the other side [38]. This might be due to other chickens perching and consequently blocking the way from one side of the pen to the other side. It might also be related to their heavy body weight, particularly in the last week of the rearing period. Whether or not a more balanced pen enrichment might have comparable stimulatory effects on activity, while maintaining performance, needs to be investigated.

Regarding the strain, in the current study, body weight and feed intake of fast-growing chickens were higher than slower-growing broiler chickens on same ages, which is in accordance with previous studies [45,66,67,68]. Due to a use of very young broiler breeders, body weight gain and BW at slaughter was relatively low [69, 70], which might have resulted in a low prevalence of leg disorders as well (see below).

### Tibia characteristics

One of the most important underlying reasons for suboptimal bone development in broiler chickens is high growth rate, while low activity levels is the other one [12,24,41]. The hypothesis of this study was that leg health and bone characteristics in broiler chickens can be improved through pen enrichment, which has previously been confirmed by several studies [21,30,31,33,35]. Focusing on bone properties, a higher activity has been found to positively affect tibia morphological, biophysical and mechanical characteristics of chickens [26,33,35,44,72,73]. A large distance between feed and water resulted in increased walking activity [18] and better tibia development in broiler chickens [26, 33]. Barrier perches resulted in improved tibia characteristics of laying hens [71]. Using sand as a dustbathing material and addition of strings for activity stimulation resulted in better bone development in fast-growing broiler chickens [72].

The results of the current study showed that tibia osseous volume, total volume and volume fraction of both fast and slower-growing broiler chickens and tibia pore volume of slower-growing chickens only were positively affected by pen enrichment, while most of the other tibia characteristics were slightly higher, but not significant. These findings are in agreement with previous studies, indicating the stimulating effects of pen enrichment on bone characteristics [33,73,74]. Tibia characteristics were found correlated with leg disorders and locomotion. Chickens with advanced tibia characteristics showed better locomotion and less leg disorders [73, 74]. In the current study, it can be suggested that bone mineral deposition is the most stimulated physiological mechanism by pen enrichment, whereas tibia morphological and mechanical characteristics, such as tibia weight, length, strength and stiffness, were not affected. It can be hypothesized that stimulated activity due to pen enrichment particularly affects physiological pathways involved in ossification and mineralization, rather than affecting anatomical and physical tibia characteristics.

Regarding the strain, almost all tibia morphological, biophysical and mechanical characteristics in both body weight classes were higher in slower-growing chickens than in fast-growing chickens. These findings are in line with previous studies, indicating that slower-growing chickens demonstrate better bone characteristics in all ages compared to fast-growing chickens [14,42,43,75,76]. Fast-growing chickens have more porous and less mineralized leg bones than slower-growing broiler chickens, which together with a higher body weight results in a higher risk of lameness [14,42,43,44,45,77], impaired activity and locomotion [24,26,46,48,49,78] and more leg problems [46, 51].

### Leg disorders and gait score

In the current study, the incidence of TD did not differ between pen enrichment, nor between strains, while other leg disorders (BCO, EPA and EPI) were hardly or not observed. These results might be explained by a relatively low stocking density (10 chickens/m^2^), which is related to a low prevalence of leg disorders [10,40,77,79]. Additionally, BW of the chickens was relatively low, probably related to the use of offspring from young broiler breeders. VV angulation in right legs was found to be higher in fast-growing chickens than in slower-growing chickens. These results are in line with previous studies, indicating that growth rate plays an important role on the prevalence of VV [12,80,81,82]. This might be explained by irregular and poor vascular morphology of the epiphyseal growth plate and insufficiently mineralized bones in fast-growing broiler chickens [81, 83]. Slower-growing chickens, on the contrary, have more time for bone mineralization, which compensates the lack of mineralization in the early growth phase, that loads less stress on the skeleton [73,83,84,85], and eventually result in a low incidence of VV. Despite the fact that VV angulation in right legs differed between strains in the current study, the maximal average angulation was 6.04° and it can be disputed whether or not this degree of angulation can be considered as VV or as a leg disorder.

Better gait was found in slower-growing chickens than in fast-growing chickens, both housed in non-enriched pens, while chickens of both strains in enriched pens had a gait score in between. It can be speculated that in fast-growing chickens provided with sufficient enrichment, locomotion can be stimulated, but in case no enrichment is present, fast-growing chickens show poorer gait scores than slower-growing chickens. Stimulating locomotion by pen enrichment might be more beneficial for fast-growing chickens, since they have less advanced bone development and poorer leg health than slower-growing chickens [38,42,74,75,85,86]. However, in the current study differences between treatments were relatively small.

### Home pen behaviour and use of enrichment

Results of home pen behaviour showed that broiler chickens in enriched pens demonstrated higher or a tendency to higher percentages of active behaviours (e.g., standing, walking, foraging) and lower percentages of passive behaviours (e.g., comfort behaviour, sitting) than chickens in non-enriched pens. This is in accordance with previous studies, indicating that pen enrichment may stimulate physical activity. Placing horizontal platforms [21,23,33], angular ramps [87] and barrier perches [30,31,32,88] resulted in stimulated activity in broiler chickens. Using wooden boxes with peat for dust bathing, two platforms with ramps and two bales of peat as a pen enrichment resulted in more wing flapping, wing stretching, body shaking, ground scratching, ground pecking and foraging behaviours in fast-growing broiler chickens compared to non-enriched pens [89]. Scattering mealworms [36] and Black Soldier fly larvae [37, 38] on the litter in fast-growing broiler chickens resulted in increased physical activity and locomotion. A large distance between feeder and drinker as a pen enrichment also resulted in a high percentage of active behaviours [38,62,90]. Different dustbathing materials, such as moss-peat have also been found to contribute to activity of broiler chickens [35, 38].

In the current study, slower-growing broiler chickens demonstrated higher or tendency to higher percentages of active behaviours (e.g., standing, walking, foraging) and lower percentages of passive behaviours (e.g., comfort behaviour, sitting) at all observation days and also on similar body weights (day 22 for fast-growing chickens and day 29 for slower-growing chickens) than fast-growing chickens. In addition, use of enrichment objects differed between fast and slower-growing broiler chickens. The most attention-grabbing difference was observed in chickens perching on barriers. A considerably higher percentage of slower-growing chickens were found perching on barriers compared to fast-growing chickens at all ages and also at the same body weight. Slower-growing chickens also showed a higher or a tendency to higher preference to go on ramps and platforms, while fast-growing chickens preferred to stay under the platforms and ramps. These findings are in in line with previous studies showing slower-growing chickens demonstrated more active behaviours than fast-growing chickens. Fast-growing broiler chickens showed higher percentages of time sitting idle and lower percentages of time standing and walking than slower-growing chickens [26,45,47,49]. Slower-growing chickens have been found to use perches more than fast-growing chickens [30,38, 91,92,93]. It has been shown that fast-growing broiler chickens showed a preference for lying and sitting on the litter instead of using raised platforms and perches. All these findings might be due to the imbalance between high growth rate and immature bones of fast-growing chickens than slower-growing chickens, which negatively affects standing, particularly at higher body weights, walking and foraging behaviours. This makes fast-growing chickens have more difficulties with barrier perches to access feed and water, to climb and go down on angular ramps than the slower-growing chickens. Another potential reason for these differences between strains might be related to body weight and heavy breast muscles. However, the current study clearly demonstrates that at the same BW class, still differences in activity related behaviours were present between the fast and slower-growing chickens, which suggests that other aspects than BW appears to play a role as well.

## Conclusion

In both slower and fast-growing chickens, tibia biophysical characteristics were positively influenced by comprehensive pen enrichment, while tibia morphological and mechanical characteristics were not affected, suggesting that pen enrichment particularly affects physiological mechanisms related to ossification and mineralization. Slower-growing chickens showed better tibia characteristics, and active behaviours than fast-growing chickens. Pen enrichment resulted in lower body weight gain in both fast and slower-growing chickens, which might be due to higher activity or lower feed intake as a result of difficulties of crossing the barrier perches. The relationship between tibia development and leg health remains unclear, because of the very low incidence of leg disorders in the current study.

## Acknowledgments

This experiment was the part of the “Healthy Bones” project, financed by a public-private partnership (TKI-AF-15203; BO-63-001-004). The financial support of the Ministry of Agriculture, Nature, and Food Quality (The Netherlands), Aviagen (UK), Darling Ingredients Inc., ForFarmers N.V., Hubbard, Marel Stork Poultry Processing BV, Nepluvi (all The Netherlands) and Trouw Nutrition (Nutreco) is gratefully acknowledged. The authors would like to thank Remco Hamoen for his expertise during the 3D micro–CT X-ray scanning. Bert van Nijhuis, Henny Reimert, Ilona van der Anker-Hensen, Bjorge Laurenssen and the animal caretakers and staff at Wageningen Bioveterinary Research (Lelystad, The Netherlands) are acknowledged for their assistance during the experiment.

## References

1. Havenstein GB, Ferket PR, Qureshi MA. Growth, livability, and feed conversion of 1957 versus 2001 broilers when fed representative 1957 and 2001 broiler diets. Poult Sci 2003; 82: 1500–1508.

2. Knowles TG, Kestin SC, Haslam SM, Brown SN, Green LE, Butterworth A, et al. Leg disorders in broiler chickens: prevalence, risk factors and prevention. PLoS OnE. 2008; 3: 1545.

3. Petracci M, Cavani C. Muscle growth and poultry meat quality issues. Nutrients. 2012; 4: 1–12.

4. Zuidhof MJ, Schneider BL, Carney VL, Korver DR, Robinson FE. Growth, efficiency, and yield of commercial broilers from 1957, 1978, and 2005. Poult Sci. 2014; 93: 2970–2982.

5. Sherlock L, Demmers TGM, Goodship AE, McCarthy ID, Wathes CM. The relationship between physical activity and leg health in the broiler chicken. Br Poult Sci. 2010; 51: 22–30.

6. González-Cerón F, Rekaya R, Aggrey SE. Genetic analysis of bone quality traits and growth in a random mating broiler population. Poult Sci. 2015; 94: 883–889.

7. Tallentire CW, Leinonen I, Kyriazakis I. Breeding for efficiency in the broiler chicken: A review. Agron Sustain Dev. 2016; 36: 1–16.

8. Gocsik É, Silvera AM, Hansson H, Saatkamp HW, Blokhuis HJ. Exploring the economic potential of reducing broiler lameness. Br Poult Sci. 2017; 58: 337–347.

9. Morris MP. National survey of leg problems. Pigs and Poultry, 1993; 6: 16.

10. Bessei W. Welfare of broilers: a review. Worlds Poult Sci J. 2006; 62: 455–466.

11. McKay JC, Barton NF, Koerhuis ANM, McAdam J. The challenge of genetic change in the broiler chicken. BSAP Occas Publ. 2000; 27: 1–7.

12. Bradshaw RH, Kirkden RD, Broom DM. A review of the aetiology and pathology of leg weakness in broilers in relation to welfare. Avian Poult Biol Rev. 2002; 13: 45–104.

13. EFSA panel on animal health and welfare. Scientific opinion on the influence of genetic parameters on the welfare and the resistance to stress of commercial broilers. EFSA journal. 2010; 8: 1666.

14. Sullivan TW. Skeletal problems in poultry: estimated annual cost and descriptions. Poult Sci. 1994; 73: 879–882.

15. Kestin SC, Su G, Sørensen P. Different commercial broiler crosses have different susceptibilities to leg weakness. Poult Sci. 1999; 78: 1085–1090.

16. Mench J. Lameness. In: Measuring and auditing broiler welfare. 2004. p. 3–17.

17. Grandin T. Auditing animal welfare at slaughter plants. Meat Sci. 2010; 86:56–65.

18. Reiter K, Bessei W. Effect of locomotor activity on leg disorder in fattening chicken. Berl Munch Tierarztl Wochenschr. 2009; 122: 264–270.

19. Blatchford RA, Archer GS, Mench JA. Contrast in light intensity, rather than day length, influences the behavior and health of broiler chickens. Poult Sci. 2012; 91: 1768–1774.

20. Ohara A, Oyakawa C, Yoshihara Y, Ninomiya S, Sato S. Effect of environmental enrichment on the behavior and welfare of Japanese broilers at a commercial farm. J Poult Sci. 2015; p. 0150034.

21. Pedersen IJ, Forkman B. Improving leg health in broiler chickens: a systematic review of the effect of environmental enrichment. Anim Welf. 2019; 28: 215–230.

22. Sandilands V, Moinard C, Sparks NHC. Providing laying hens with perches: fulfilling behavioural needs but causing injury?. Brit Poult Sci. 2009; 50: 395–406.

23. Kaukonen E, Norring M, Valros A. Effect of litter quality on foot pad dermatitis, hock burns and breast blisters in broiler breeders during the production period. Avian Pathol. 2016; 45: 667–673.

24. Weeks CA, Danbury TD, Davies HC, Hunt P, Kestin SC. The behaviour of broiler chickens and its modification by lameness. Appl Anim Behav Sci. 2000; 67: 111–125.

25. Reiter K, Bessei W. Possibilities to reduce leg disorders in broilers and turkeys. Arch Für Geflügelkunde. 1998; 62: 145–149.

26. Reiter K, Bessei W. Effect of locomotor activity on bone development and leg disorders in broilers. Arch Für Geflügelkunde (Germany). 1998.

27. Balog JM, Bayyari GR, Rath NC, Huff WE, Anthony NB. Effect of intermittent activity on broiler production parameters. Poult Sci. 1997; 76: 6–12.

28. Martrenchar A, Huonnic D, Cotte JP, Boilletot E, Morisse JP. Influence of stocking density, artificial dusk and group size on the perching behaviour of broilers. Br Poult Sci. 2000; 41: 125–130.

29. Hall AL. The effect of stocking density on the welfare and behaviour of broiler chickens reared commercially. Anim Welf. 2001; 10: 23–40.

30. Ventura, B. A., F. Siewerdt, and I. Estevez. 2012. Access to barrier perches improves behavior repertoire in broilers. PloS one. 7:29826.

31. Zhao JP, Jiao HC, Jiang YB, Song ZG, Wang XJ, Lin H. Cool perches improve the growth perfor-mance and welfare status of broiler chickens reared at different stocking densities and high temperatures. Poult Sci. 2013; 92: 1962–1971.

32. Kaukonen E, Norring M, Valros A. Perches and elevated platforms in commercial broiler farms: use and effect on walking ability, incidence of tibial dyschondroplasia and bone mineral content. Animal. 2017; 11: 864–871.

33. Pedersen IJ, Tahamtani FM, Forkman B, Young JF, Poulsen, HD, Riber AB. Effects of environmental enrichment on health and bone characteristics of fast growing broiler chickens. Poult Sci. 2020; 99: 1946–1955.

34. Bizeray D, Estevez I, Leterrier C, Faure JM. Effects of increasing environmental complexity on the physical activity of broiler chickens. Appl Anim Behav Sci. 2002; 79: 27–41.

35. Riber AB, van de Weerd HA, De Jong IC, Steenfeldt S. Review of environmental enrichment for broiler chickens. Poult Sci. 2018; 97: 378–396.

36. Pichova K, Nordgreen J, Leterrier C, Kostal L, Moe RO. The effects of food-related environmental com-plexity on litter directed behaviour, fear and exploration of novel stimuli in young broiler chickens. Appl Anim Behav Sci. 2016; 174: 83–89.

37. Ipema AF, Gerrits WJ, Bokkers EA, Kemp B, Bolhuis JE. Provisioning of live black soldier fly larvae (Hermetia illucens) benefits broiler activity and leg health in a frequency-and dose-dependent manner. Appl Anim Behav Sci. 2020; 230: p. 105082.

38. De Jong IC, Blaauw XE, Van der Eijk JAJ, Da Silva CS, Van Krimpen MM, et al. Providing environmental enrichments affects activity and performance, but not leg health in fast- and slower-growing broiler chickens. Appl Anim Behav Sci. 2021; 105375.

39. Yıldız H, Petek M, Sönmez G, Arıcan I, Yılmaz B. Effects of lighting schedule and ascorbic acid on performance and tibiotarsus bone characteristics in broilers. Turkish J Vet Anim Sci. 2009; 33: 469–476.

40. Buijs S, van Poucke E, Van Dongen S, Lens L, Baert J, Tuyttens FA. The influence of stocking density on broiler chicken bone quality and fluctuating asymmetry. Poult Sci. 2012; 91: 1759–1767.

41. Van der Pol CW, Molenaar R, Buitink CJ, Van Roovert-Reijrink IAM, Maatjens CM, van den Brand H, et al. Lighting schedule and dimming period in early life: consequences for broiler chicken leg bone development. Poult Sci. 2015; 94: 2980–2988.

42. Williams B, Solomon S, Waddington D, Thorp B, Farquharson C. Skeletal development in the meat-type chicken. Br Poult Sci. 2000; 41: 141–149.

43. Torres CA, Korver DR. Influences of trace mineral nutrition and maternal flock age on broiler embryo bone development. Poult Sci. 2018; 97: 2996–3003.

44. Williams B, Waddington D, Murray DH, Farquharson C. Bone strength during growth: influence of growth rate on cortical porosity and mineralization. Calcif Tissue Int. 2004; 74: 236–245.

45. Stojcic MD, Bessei W. The effect of locomotor activity and weight load on bone problems in fast and slow growing chickens. Arch für Geflügelkunde. 2009; 73: 242–249.

46. Bokkers EA, Koene P. Motivation and ability to walk for a food reward in fast and slower-growing broilers to 12 weeks of age. Behav Process. 2004; 67: 121–130.

47. Wallenbeck A, Wilhelmsson S, Jönsson L, Gunnarsson S, Yngvesson J. Behaviour in one fast growing and one slow-growing broiler (Gallus gallus domesticus) hybrid fed a high-or low-protein diet during a 10-week rearing period. Acta Agr Scand A—An Sci. 2016; 66: 168–176.

48. Lewis PD, Perry GC, Farmer LJ, Patterson RLS. Responses of two genotypes of chicken to the diets and stocking densities typical of UK and ‘Label Rouge production systems: I. Performance, behaviour and carcass composition. Meat Sci. 1997; 45: 501–516.

49. Cornetto T, Estevez I. Behavior of the domestic fowl in the presence of vertical panels. Poult Sci. 2001; 80: 1455–1462.

50. Reiter K, Kutritz B. Behaviour and leg weakness in different broiler breeds. Arch für Geflügelkunde. 2001; 65: 137–141.

51. Kjaer JB, Su G, Nielsen BL, Sørensen P. Foot pad dermatitis and hock burn in broiler chickens and degree of inheritance. Poult Sci. 2006; 85: 1342–1348.

52. Havenstein GB, Ferket PR, Scheideler SE, Larson BT. Growth, livability, and feed conversion of 1957 vs 1991 broilers when fed “typical” 1957 and 1991 broiler diets. Poult Sci. 1994; 73: 1785–1794.

53. De Jong IC, Gunnink H. Effects of a commercial broiler enrichment programme with or without natural light on behaviour and other welfare indicators. Anim. 2019; 13: 384–391.

54. Kestin SC, Knowles TG, Tinch AE, Gregory NG. Prevalence of leg weakness in broiler chickens and its relationship with genotype. Vet Rec. 1992; 131: 190–194.

55. Güz BC, Molenaar R, De Jong IC, Kemp B, Van Krimpen M, Van Den Brand H. Effects of eggshell temperature pattern during incubation on tibia characteristics of broiler chickens at slaughter age. Poult Sci. 2020; 99: 3020–3029.

56. Güz BC, Molenaar R, De Jong IC, Kemp B, Van Krimpen M, Van den Brand H. Effects of green light emitting diode light during incubation and dietary organic macro and trace minerals during rearing on tibia characteristics of broiler chickens at slaughter age. Poult Sci. 2021; 100: 707–720.

57. Riesenfeld A. Metatarsal robusticity in bipedal rats. Am J Phys Anthropol. 1972; 36: 229–233.

58. Jungmann R, Schitter G, Fantner GE, Lauer ME, Hansma PK, Thurner PJ. Real-time microdamage and strain detection during micromechanical testing of single trabeculae. In: Experimental and Applied Mechanics: SEM Annual Conference and Exposition. 2007. pp. 0–11.

59. Novitskaya E, Chen PY, Hamed E, Jun L, Lubarda VA, Jasiuk I, et al. Recent advances on the measurement and calculation of the elastic moduli of cortical and trabecular bone: a review. Theor Appl Mech. 2011; 38: 209–297.

60. Turner CH, Burr DB. Basic biomechanical measurements of bone: a tutorial. Bone. 1993; 14: 595–608.

61. Bach MH, Tahamtani FM, Pedersen IJ, Riber AB. Effects of environmental complexity on behaviour in fast-growing broiler chickens. Appl Anim Behav Sci. 2019; 219: pp. 104840.

62. Jones PJ, Tahamtani, FM, Pedersen IJ, Niemi JK, Riber AB. The productivity and financial impacts of eight types of environmental enrichment for broiler chickens. Animals. 2020; 10: 378.

63. Pettit-Riley R, Estevez I. Effects of density on perching behavior of broiler chickens. Appl Anim Behav Sci. 2001; 71: 127–140.

64. Simsek UG, Dalkilic B, Ciftci M, Cerci IH, Bahsi M. Effects of enriched housing design on broiler performance, welfare, chicken meat composition and serum cholesterol. Acta Vet Brno. 2009; 78: 67–74.

65. Yildirim M, Taskin A. The effects of environmental enrichment on some physiological and behavioral parameters of broiler chicks. Braz J Poult Sci. 2017; 19: 355–362.

66. Benyi K, Acheampong-Boateng O, Norris D, Ligaraba TJ. Response of Ross 308 and Hubbard broiler chickens to feed removal for different durations during the day. Trop Anim Health Pro. 2010; 42: 1421–1426.

67. Benyi K, Netshipale AJ, Mahlako KT, Gwata ET. Effect of genotype and stocking density on broiler performance during two subtropical seasons. Trop Anim Health Pro. 2015; 47: 969–974.

68. Dixon LM. Slow and steady wins the race: The behaviour and welfare of commercial faster growing broiler breeds compared to a commercial slower growing breed. PLoS ONE. 2020; 15: e0231006.

69. Peebles ED, Doyle SM, Pansky T, Gerard PD, Latour MA, Boyle CR, et al. Effects of breeder age and dietary fat on subsequent broiler performance. 1. Growth, mortality, and feed conversion. Poult Sci. 1999; 78: 505–511.

70. Nasri H, Van den Brand H, Najjar T, Bouzouaia M. Interactions between egg storage duration and broiler breeder age on egg fat content, chicken organ weights, and growth performance. Poult Sci. 2020; 99: 4607–4615.

71. Hughes BO, Wilson S, Appleby MC, Smith SF. Comparison of bone volume and strength as measures of skeletal integrity in caged laying hens with access to perches. Res Vet Sci. 1993; 54: 202–206.

72. Arnould C, Bizeray D, Faure JM, Leterrier C. Effects of the addition of sand and string to pens on use of space, activity, tarsal angulations and bone composition in broiler chickens. Anim Welf. 2004; 13: 87–94.

73. Toscano MJ, Nasr MAF, Hothersall B. Correlation between broiler lameness and anatomical measurements of bone using radiographical projections with assessments of consistency across and within radiographs. Poult Sci. 2013; 92: 2251–2258.

74. Sørensen P, Su G, Kestin SC. Effects of age and stocking density on leg weakness in broiler chickens. Poult Sci. 2000; 79: 864–870.

75. Lilburn MS. Skeletal growth of commercial poultry species. Poult Sci. 1994; 73: 897–903.

76. Thorp BH, Waddington D. Relationships between the bone pathologies, ash and mineral content of long bones in 35-day-old broiler chickens. Res Vet Sci. 1997; 62: 67–73.

77. Shim MY, Karnuah AB, Mitchell AD, Anthony NB, Pesti GM, Aggrey SE. The effects of growth rate on leg morphology and tibia breaking strength, mineral density, mineral content, and bone ash in broilers. Poult Sci. 2012; 91: 1790–1795.

78. Siegel PB, Honaker CF, Rauw WM. Selection for high production in poultry. CABI Publishing: Wallingford, UK. 2009; pp. 230–242.

79. Buijs S, Keeling L, Rettenbacher S, Van Poucke E, Tuyttens FAM. Stocking density effects on broiler welfare: Identifying sensitive ranges for different indicators. Poult Sci. 2009; 88: 1536–1543.

80. Leterrier C, Nys Y. Clinical and anatomical differences in varus and valgus deformities of chick limbs suggest different aetio-pathogenesis. Avian Pathol. 1992; 21: 429–442.

81. Shim MY, Karnuah AB, Anthony NB, Pesti GM, Aggrey SE. The effects of broiler chicken growth rate on valgus, varus, and tibial dyschondroplasia. Poult Sci. 2012; 91: 62–65.

82. González-Cerón F, Rekaya R, Anthony NB, Aggrey SE. Genetic analysis of leg problems and growth in a random mating broiler population. Poult Sci. 2015; 94: 162–168.

83. Kestin SC, Gordon S, Su G, Sørensen P. Relationships in broiler chickens between lameness, liveweight, growth rate and age. Vet Rec. 2001; 148: 195–197.

84. Thorp BH. Skeletal disorders in the fowl: a review. Avian Path. 1994; 23: 203–236.

85. Sanchez-Rodriguez E, Benavides-Reyes C, Torres C, Dominguez-Gasca N, Garcia-Ruiz AI, Gonzalez-Lopez S, et al. Changes with age (from 0 to 37 D) in tibiae bone mineralization, chemical composition and structural organization in broiler chickens. Poult Sci. 2019; 98: 5215–5225.

86. Rayner AC, Newberry RC, Vas J, Mullan S. Slow-growing broilers are healthier and express more behavioural indicators of positive welfare. Sci Rep. 2020; 10: 1–14.

87. Birgul OB. Mutaf S, Alkan S. Effects of different angled perches on leg disorders in broilers. Arch für Geflügelkunde. 2012; 76: 44–48.

88. Bench CJ, Oryschak MA, Korver DR, Beltranena E. Behaviour, growth performance, foot pad quality, bone density, and carcass traits of broiler chickens reared with barrier perches and fed different dietary crude protein levels. Can J Anim Sci. 2016; 97: 268–280.

89. Vasdal G, Vas J, Newberry RC, Moe RO. Effects of environmental enrichment on activity and lameness in commercial broiler production. J Appl Anim Welf Sci. 2019; 22: 197–205.

90. Reiter K, Bessei W. Effect of the distance between feeder and drinker on behaviour and leg disorders of broilers. In: Proceedings of the 30th International Congress of the International Society for Applied Ethology. 1996.

91. Groves PJ, Muir WI. June. Use of perches by broiler chickens in floor pen experiments. In: Proceedings of the IX European Symposium on Poultry Welfare. 2013. pp. 17-20.

92. Yngvesson J, Wedin M, Gunnarsson S, Jönsson L, Blokhuis H, Wallenbeck A. Let me sleep! Welfare of broilers (Gallus gallus domesticus) with disrupted resting behaviour. Acta Agr Scand A—An Sci. 2017; 67: 123–133.

93. Bailie CL, O’Connell NE. Perch design preferences of commercial broiler chickens reared in windowed houses. In Proc. EAAP 67th Annual Meeting. 2016ç

